# Effects of sex and pre-exposure on Δ9-tetrahydrocannabinol (THC) vapor self-administration in rats

**DOI:** 10.1101/2025.08.01.668172

**Authors:** Catherine F. Moore, Elise M. Weerts

## Abstract

**Rationale:** Animal models of cannabinoid self-administration are critical for advancing our understanding of the neurobiology of cannabis use and for developing medications for Cannabis Use Disorder. Use of vapor inhalation models have translational relevance, as the majority of human cannabis user do so by inhalation (e.g., smoking, vaporization).

**Methods:** Adult male and female Sprague Dawley rats (N=96, 6-12 per sex/group) were pre-exposed to vaporized THC or vehicle (Veh; 100% propylene glycol). Rats were then trained to self-administer vapor puffs of either THC (50 mg/ml training dose) or Veh under a fixed ratio (FR) 1 schedule. Responding was then assessed under increasing response costs (FR1-5) and different ‘doses’ (50-200 mg/ml) of THC. As a secondary study aim, we assessed the effects of pre-exposure to THC (or Veh) on self-administration of THC vapor or Veh vapor.

**Results:** There were no differences in responding for THC and Veh vapor under an FR1 schedule. As the FR increased, rats increased their responses for THC, and female rats in the THC group responded more than female rats in the Veh vapor group under FR4-5 schedules. When THC concentration was varied, rats titrated their intake in a predictable U-shaped pattern. THC vapor pre-exposure significantly increased self-administration of THC vapor, but not Veh vapor.

**Conclusions:** Male and female rats voluntarily administered THC vapor in a sustained manner across several months, replicating what has been shown with vaporized cannabis extracts male rats. This study is the first to demonstrate that THC vapor has reinforcing properties in female rats and that pre-exposure is critical for engendering higher intake. This is a promising model of voluntary THC vapor administration.

## 1. Introduction

Cannabis use is widespread and increasing in the United States. A significant portion of individuals who use cannabis go on to develop Cannabis Use Disorder (CUD) and/or experience problematic cannabis use, which is a subclinical threshold of use that leads to negative health and/or social consequences. Preclinical models of drug self-administration are useful for assessing drug abuse liability, as well as the behavioral and neurobiological correlates of problematic drug use and ‘addiction-like’ phenotypes (Deroche-Gamonet 2020). Self-administration refers to an operant procedure where a trained behavioral response (e.g., nose poke) is maintained by drug delivery. Reinforcement is demonstrated if the behavioral response associated with drug delivery exceeds responding for the drug vehicle. Self-administration procedures also provide a model to evaluate the effects of voluntary intake.

There have been many challenges in establishing preclinical models of cannabinoid self-administration, particularly for delta-9-tetrahydrocannabinol (THC) (Tanda 2016). Standard preclinical drug self-administration procedures use the intravenous (IV) route, due to its rapid onset of effects and precise drug delivery. Early studies investigating IV self-administration of THC in rodents did not show levels of responding higher than vehicle (Takahashi and Singer 1979, 1980; van Ree et al. 1978). However, selective high affinity CB1R agonists (e.g., WIN 55,212-2), which have greater potency than THC, are self-administered in a dose-dependent manner by rodents (Martellotta et al. 1998; Fattore et al. 2001; Deiana et al. 2007; Fattore, Spano, et al. 2007). Though even in rats trained to self-administer WIN 55,212-2, substitution of THC did not maintain responding (Lefever et al. 2014). THC was demonstrated to have reinforcing effects when administered IV in squirrel monkeys (Justinova et al. 2003; Tanda et al. 2000). Success of this group was attributed to the use of very low THC doses (2-4 µg/kg/injection), use of a novel vehicle to enhance solubility of THC, rapid drug delivery, and exposure to full range of doses (Panlilio et al. 2010).

Several factors contribute to the difficulty of establishing IV THC self-administration in rodents. IV catheters require maintenance, as patency issues can occur and limit their functional life. Additionally, low solubility of THC for IV delivery necessitates higher volumes and slower injection speeds, which can affect both pharmacokinetics and reinforcement. Importantly, at higher doses, THC can also produce aversive effects, which may underlie some of the challenges with developing self-administration models. However, tolerance develops to the initial aversive effects of THC, and studies using place conditioning have demonstrated that pre-exposure to THC can attenuate its unconditioned aversive effects (Murray and Bevins 2010).

In recent years, there have been many advances in preclinical inhalation models for cannabis self-administration. These models have good translational relevance, as cannabis is primarily used by humans via inhalation smoked and vaporized delivery methods (Morean et al. 2017; Giroud et al. 2015; Budney et al. 2015). Vapor procedures are non-invasive and can be maintained over longer periods (months-years) allowing systematic within subject manipulations of drug dose, access conditions, and duration of exposure. Freels et al. (2020) recently demonstrated that vaporized cannabis extract, rich in delta-9-tetrahydrocannabinol (THC), was self-administered by male rats at rates higher than vehicle vapor or a vaporized cannabidiol (CBD)-rich extract under fixed-ratio (FR) 4 and progressive ratio schedules. Subsequent studies have shown that both male and female rats will voluntarily administer cannabis vapor, with delivery-dependent increases in drug plasma levels as well as behavioral and biological changes in cannabis vapor administering groups (Glodosky et al. 2020; Freels et al. 2024). However, in these studies, animals responded for vapor under an FR1 schedule, and the levels of responding for THC and vehicle vapor were similar under these parameters. These studies have all used cannabis extracts that contains various phytocannabinoids and terpenes found in cannabis.

In the present study, we wanted to establish whether male and female rats would self-administer vaporized THC isolate over vaporized vehicle. To increase our likelihood of success, we leveraged several design features which have been shown to augment taking of drugs that may have sedative effects and utilized species-specific features. First, sessions were run during the dark cycle when rats are most active, similar to ‘drinking in the dark’ procedure where alcohol access occurs in the dark cycle, which is a highly successful model of alcohol binge drinking (Holgate et al. 2017). Second, we tested intermittent access procedures, where self-administration days alternate with abstinence days. Intermittent access has been show to produce an escalation of drug self-administration and an increase in behavioral sensitization for other drugs of abuse (O’Dell and Koob 2007; Mandyam et al. 2008; Calipari et al. 2015; Kawa et al. 2016; Carnicella et al. 2014; Goodwin 2016). Another key factor is use of environmental stimuli (e.g., cues) that are at first neutral with respect to a particular behavior and then repeatedly paired with a drug and can acquire positive or negative incentive values. These values are defined in terms of the behavioral responses produced by the stimulus (e.g., exploration and approach vs. avoidance and escape). It is well established that drug-related stimuli and associated “cue reactivity” (Wikler 1973) can play an important role in the maintenance of drug seeking and self-administration (Fattore, Fadda, et al. 2007; Justinova et al. 2008; Markou et al. 2016; Kenny et al. 2006; Markou et al. 1993; Arroyo et al. 1998). Thus, our setup included distinct stimuli to facilitate and maintain self-administration. We included a pre-exposure component (non-contingent), prior to contingent drug delivery, as data from CPP studies show that pre-exposure to THC can attenuate its unconditioned aversive effects (Murray and Bevins 2010). Finally, we selected the THC vapor concentration for self-administration (50 mg/ml) based on our own preliminary data and to match Freels et al. (2020) cannabis extract’s THC content.

All rats received pre-exposure to their respective self-administration drug (THC or Veh) prior to self-administration training as part of a separate study on the conditioned rewarding effects of vaporized THC, results which are published elsewhere (Moore et al. 2024). In these animals, we assessed operant responding for deliveries of a training ‘dose’ of 50 mg/ml THC vapor under an FR1 schedule, and then evaluated the impact of increased response costs, and ‘dose’-effects on THC vapor self-administration behavior. As a secondary study aim, we assessed the effects of pre-exposure to THC (or Veh) on self-administration of THC vapor or Veh vapor in male and female rats.

## 2. Materials and Methods

### 2.1 Subjects

Adult male and female Sprague Dawley rats (N=96, 6-12 per sex/group)(Charles River, Wilmington, MA), 8 weeks old at the start of experiments, were single housed in wire-topped, plastic cages (27 × 48 × 20 cm) with standard enrichment. The vivarium was on a 12 hr reverse light cycle (lights off at 8:00 a.m.) and was humidity and temperature controlled. Diet was a corn-based chow (Teklad Diet 2018; Harlan, Indianapolis, IN); rats were food maintained to 90% of their free-feeding weight throughout the duration of the experiments. Food was given daily at the end of their drug exposure/self-administration sessions. Rats had *ad libitum* access to water except during test procedures. All procedures used in this study were approved by the Johns Hopkins Institutional Animal Care and Use Committee. The facilities adhered to the National Institutes of Health *Guide for the Care and Use of Laboratory Animals* and were AAALAC-approved.

### 2.2 Drugs

THC stock solutions (50-200 mg/mL in 95% ethanol), confirmed as >95% purity, were provided by the U.S. National Institute on Drug Abuse Drug Supply Program. Ethanol from the THC stock was evaporated using nitrogen and then THC was mixed in 100% propylene glycol to yield desired mg/ml THC concentration for vaporization.

### 2.3 Vapor exposure systems

Two different commercial vapor chamber systems were used in the course of this study, both manufactured by La Jolla Alcohol Research Institute (La Jolla CA). The first was a passive exposure system which contained four sealed polycarbonate rat cages (13.5 × 11 × 10 in) adapted for the delivery of vaporized drug via an electronic vapor device (Smok Baby Beast Brother TFV8 Sub‐Ohm Tank with the V8 X‐Baby M2 0.25‐Ω coil; SMOKTech, Shenzhen, China) and connected to an air pump to regulate airflow. The chamber air was vacuum controlled by a pump which pulls room ambient air into the chamber through an intake valve and out through the exhaust valve at a constant rate (1 L/min). The evape controller was set to a maximum temperature of 400°F (∼30W), which is the same temperature used for vaporization in our clinical research (Spindle et al. 2020). The second system was set up to the mirror the parameters of the passive system, but each sealed polycarbonate chamber (13.5 × 9.0 × 8.25 in) was outfitted with 3 illuminated nose pokes across the back wall, 2 LED stimulus lights above the right and left nose poke, and 2 additional LED stimulus lights on the left and right wall above a vapor port inlet. Each chamber had its own vapor tank and device, set to 400°F maximum temperature, with chamber airflow regulated to 1 L/min. To measure amounts of e-liquid vaporized, tanks were weighed at the start and end of each day and an estimated mL per puff was calculated.

### 2.4 Vapor exposure prior to self-administration training

Rats were passively exposed to fixed amounts of THC and/or VEH vapor in 16 daily 30-min sessions, for place conditioning. Vehicle vapor was 100% PG, THC conditions were 5 puffs of 100 mg/ml (low), 5 puffs of 200 mg/ml (medium), or 10 puffs of 200 mg/ml (high) THC. Detailed methods and results from this portion of the study have been published previously (Moore et al. 2024). After a 1-month washout period, operant training was initiated. Only rats from the from the vehicle, medium THC, and high THC pre-exposure groups were used for self-administration (N=72). Rats were allocated to vehicle and THC self-administration groups and were counterbalanced based on their pre-exposure history (See Table 1).

**Table 1.**
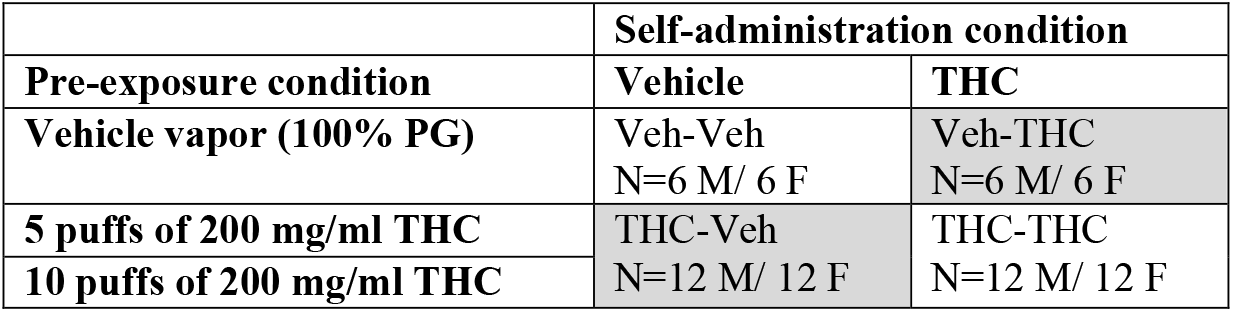
White shaded groups represent ‘matched’ pre-exposure and self-administration groups; data from these animals were analyzed for self-administration of THC compared with Veh vapor. Gray shaded groups represent ‘mismatched’ pre-exposure and self-administration groups; data from these animals were analyzed separately to evaluate how pre-exposure affects self-administration of either Veh or THC vapor.

### 2.5 Operant training of food self-administration

Rats were trained to respond for food pellets in experimental chambers (Med Associates, St. Albans, VT) equipped with an illuminated nose-poke with a response key, and a food cup connected to an automated pellet feeder, all positioned inside a sound-attenuating enclosure with exhaust fans. Rats were initially trained to press a nose poke key for delivery of a 45 mg food pellet (grain-based chow #F0165, BioServ, Flemington, NJ) in daily sessions (Mon-Fri). The nose poke key was positioned on either the left or the right side of the chamber, matched for the side of the chamber of their ‘active’ vapor lever during vapor self-administration. There was a time-out of 2.5 seconds after each pellet delivery, and sessions ended after 30-min or a maximum of 200 pellets. Initially, responding was reinforced under a fixed-ratio (FR) 1 schedule, and the FR requirement was gradually increased by 1 to a final FR10 over the course of 5 sessions. The FR was increased during the sessions based on individual performance (i.e., responding was maintained and did not decrease over consecutive reinforcers).

### 2.6 Vapor Self-Administration

Following operant food self-administration training, all self-administration sessions were conducted in the vapor chambers, and each nosepoke response resulted in delivery of vaporized THC (50 mg/ml) or vehicle (100% PG) under an FR1 schedule. Session were conducted on intermittent days, 3 times a week (e.g., Monday-Wednesday-Friday) to mirror early patterns of cannabis use in humans (Passarotti et al. 2015). At the start of each session, the houselight and the active nosepoke were illuminated. A response on the vapor-associated nosepoke resulted in a 3s vapor delivery, illumination of a cue-light on the side wall, and an extinguishing of the houselight. There was a 30-second timeout after each vapor delivery, during which the vapor associated cue light stayed on and the houselight stayed off. Two inactive nosepokes (middle and side) were never illuminated and responses were recorded but had no consequence. Sessions ended after 60-min.

The FR1 schedule continued for at least 15 sessions and until responding was stable with no increasing or decreasing trend at a group level. The FR was then progressively increased by 1 and then progressed to the next FR with a final schedule of FR5, using the following criteria: Each FR schedule was maintained for at least 5 sessions, or responding was stable with no increasing or decreasing trends before progressing to the next FR condition. Next, after stabilization of responding under the THC training dose condition (50 mg/ml) the concentration of THC in the e-liquid was increased (100-200 mg/ml) or decreased (5-10 mg/ml) in a semi-randomized order; all other variables were held constant. Rats were maintained on each concentration for at least 5 sessions and until responding was stable with no increasing or decreasing trends, before proceeding to the next dose condition.

### 2.7 Data Analysis

Self-administration data were analyzed for rats whose self-administration condition was the same as their pre-exposure condition (i.e., Veh-Veh or THC-THC), so all rats self-administering vehicle vapor had never been exposed to THC and all the rats self-administering THC vapor had been previously exposed to THC. Outcome variables were operant responses on active and inactive levers and number of vapor deliveries. For FR1 data, deliveries, active responses, and inactive responses were analyzed with a three-way ANOVA with between subjects effects of group (Veh, THC) and sex (M, F), and within subjects effects of session. To assess self-administration under increasing FR schedules, deliveries, active responses, and inactive responses were averaged across the final 3 sessions of each FR and then analyzed using a three-way ANOVA with effects of group (Veh, THC), sex (M, F) and FR (1-5). To assess self-administration across various THC concentrations, deliveries, active responses, and inactive responses were averaged across the final 3 sessions of each concentration and then analyzed using a three-way ANOVA with effects of group (Veh, THC), sex (M, F) and concentration (5-200 mg/ml). As sex differences were an a priori interest of the study, for each outcome, we also ran separate ANOVAs in males and females to conduct post-hoc testing. Dunnett’s post-hoc tests were run on within subjects factors (e.g., FR, concentration) to compare against the training conditions, and Fisher’s LSD post-hoc tests were used to compare groups. One male rat in the Veh group accidentally received THC during the 15^th^ session of FR1 self-administration testing; his data are therefore included in FR1 analysis (with data from his 15^th^ and final session being an average of the prior 3 sessions) but was then removed from the study following the exposure to THC vapor.

To evaluate the effect of pre-exposure condition on self-administration behavior, we compared rats with different pre-exposure conditions (Veh or THC) on later vapor self-administration (Veh or THC). We compared Veh-THC vs. THC-THC and Veh-Veh groups vs. THC-Veh groups using the same ANOVAs as described above, with ‘group’ (Veh, THC) now referring to pre-exposure condition.

Statistics were performed in GraphPad Prism 9 (GraphPad Software, San Diego, CA) with p ≤ 0.05 for significance. The data were Greenhouse-Geisser corrected where Mauchly’s Sphericity tests were significant.

## 3. Results

### 3.1 Active responding under an FR1 schedule

In rats self-administering Veh or THC that had pre-exposure to the same vapor condition (i.e., THC-pre-exposed self-administering THC vapor or Vehicle-pre-exposed self-administering Vehicle vapor; Veh: N=6 males, 6 females; THC: N=12 males, 12 females), males showed stable responding and completed the FR1 phase after 15 sessions. In contrast, females exhibited a slight decreasing response trend from session 13-15, so their data include an additional 5 sessions under the FR1 schedule until stability, for a total of 20 sessions. Analysis of FR1 acquisition below includes the first 15 sessions.

Under an FR1 schedule of reinforcement, there were similar amounts of responding in rats of both sexes and drug groups. There was no effect of sex or THC on number of active responses/number of b puffs obtained (under an FR1 schedule, these measures are equal). There was a main effect of session (F(5.5, 176.2) = 3.50, p<0.003) (Fig. 1a-b). ANOVAs run separately for males and females also only had significant main effects of session and not drug group. To evaluate effects across sessions, average deliveries from the first 3 and last 3 sessions were compared. Similarly, in this ANOVA, there was also only a main effect of session (F(1, 32) = 22.51, p<0.001), with post-hoc tests showing no differences between any of the 4 groups in deliveries in the first 3 or last 3 sessions. Collapsing across sex, both groups (Veh and THC) self-administered less puffs in the final 3 sessions compared with the first 3 (p’s<0.01) (Fig. 1c).

**Fig. 1.**
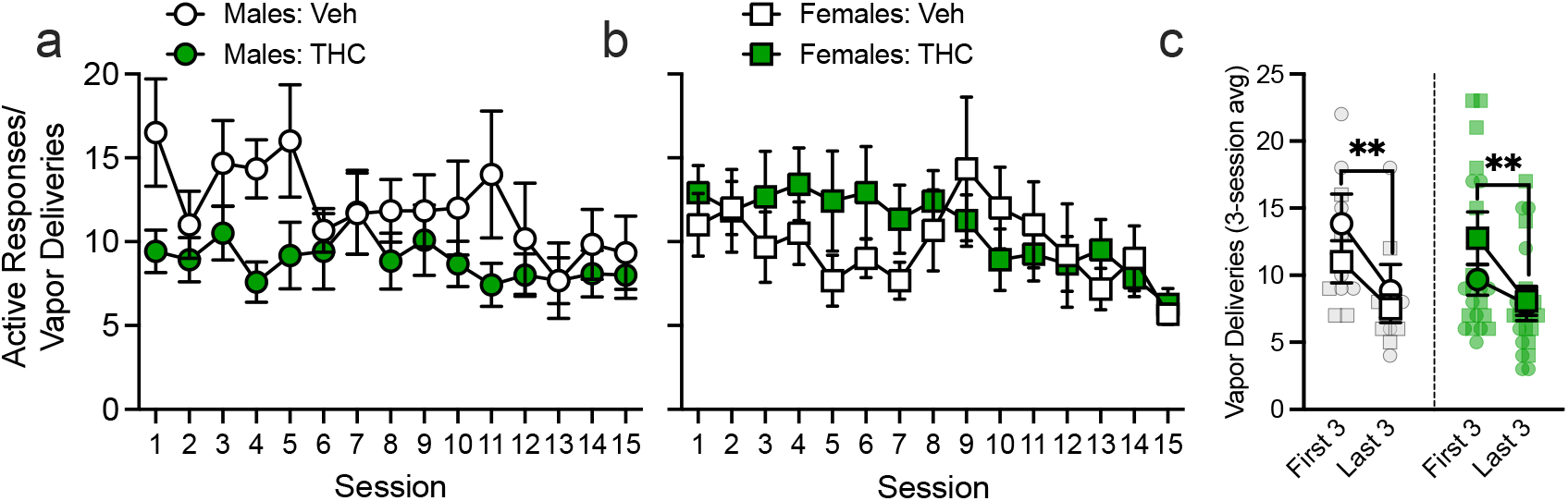
Active responding for THC or Veh vapor under an FR1 schedule in male (a) and female (b) rats. Responses decreased from the first 3 sessions to the last 3 sessions (c). Data are Mean± SEM; N=6/sex in vehicle group (white symbols), N=12/sex in THC group (green symbols). Males shown in circle symbols and females shown in square symbols. ^**^ denotes p<0.01

### 3.2 Active responding under increasing FR (1-5) schedules

As the FR requirement was increased from 1-5, responses increased across all groups to meet the higher requirements; females self-administering THC had higher responding than females self-administering Veh under FR4 and FR5 (Fig. 2a-b). In the 3-way ANOVA, there was an FR x Sex x THC interaction (F (4, 124) = 3.42, p<0.01). In an ANOVA of males only, both groups increased their responding as the FR requirement increased; responding was higher in the Veh group compared with THC under the FR3 schedule. In an ANOVA of females only, the THC group increased responding across increasing FRs (p’s<0.05 vs. FR1), while the Veh group did not increase the number of responses made (p>0.05 vs FR1). Further, females in the THC group had higher responding than the Veh group under higher FR requirements (FR4, p=0.08, FR5 (p<0.05).

**Fig. 2.**
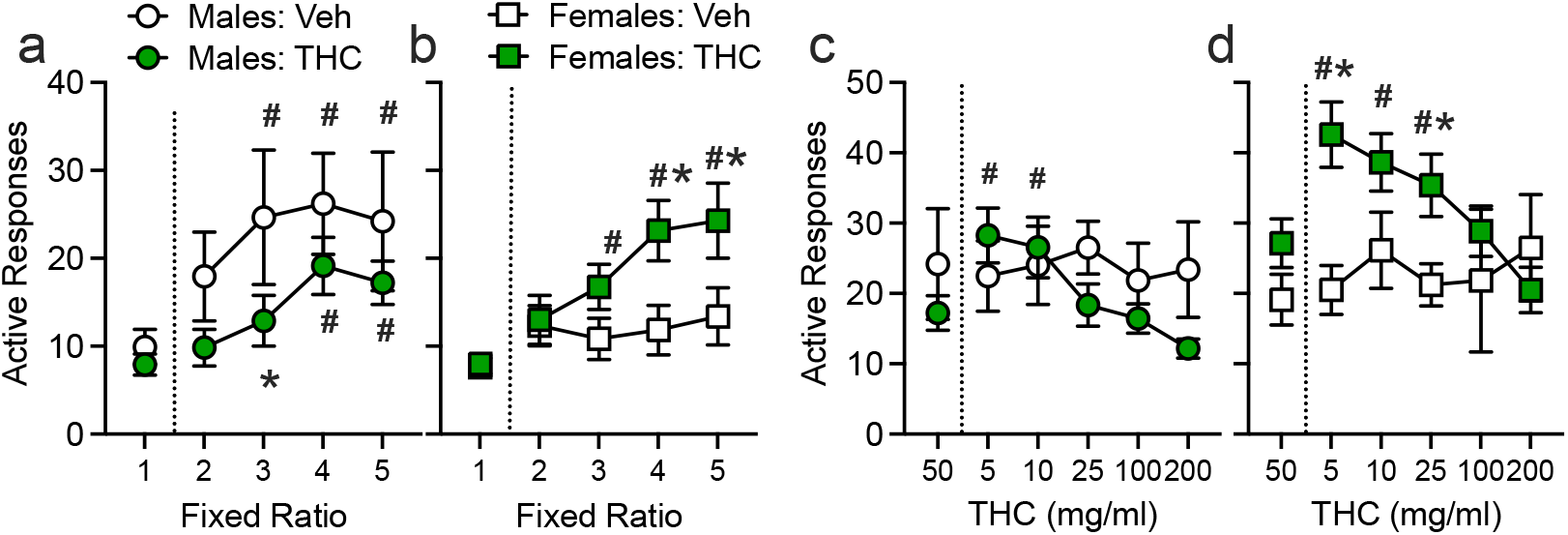
Active responding for THC or Veh vapor under increasing FR schedules in male (a) and female (b) rats. Active responding for THC or Veh at different THC concentrations (and matched Veh sessions) in male (b) and female (d) rats. Data are Mean± SEM. N=5-6/sex in vehicle groups (white symbols), N=12/sex in THC groups (green symbols). Males shown in circle symbols and females shown in square symbols. ^*^ denotes p<0.05 vs. Veh vapor group; # denotes a difference from FR1 or 50 mg/ml (training conditions)

### 3.3 Active responding for varying THC concentrations (FR5 schedule)

For rats in the THC group, as the THC concentration was decreased (i.e., lower dose), rats increased their responding to administer more puffs and as the THC concentration increased, rats decreased their responding to administer fewer puffs (Fig. 2c-d). There was a concentration x THC interaction (F (5, 155) = 7.84, p<0.0001), and no main effect nor interactions with sex. To interrogate this interaction, we conducted a two-way ANOVA with sexes collapsed; post-hoc analyses show higher responding at 5 and 10 mg/ml vs. the 50 mg/ml training concentration (p’s<0.05) and lower responding at the 200 mg/ml (p<0.05). When 5 mg/ml THC was available, the THC group had higher responding than the Veh group (p<0.05), and when 200 mg/ml THC was available, the THC group tended to have fewer responses than the Veh group (p=0.08). When analyzed within each sex separately, results indicate that females were driving this high responding: females self-administering THC had higher responding at lower THC concentrations as compared to the Veh group (5 and 25 mg/ml, p’s<0.05) and compared to the training dose (5-25 mg/ml, p’s<0.05). Responding by the males in the THC group was not different from the Veh group, but was lower at the higher THC concentrations compared to the training dose (100-200 vs. 50 mg/ml, p’s<0.05)

### 3.4 Inactive responding

Inactive responding during acquisition was high, particularly in the THC group, and this decreased across the sessions (Fig. 3). In an ANOVA of inactive responses, there was a main effect of THC (F(1, 32) = 9.78, p<0.05), but no effects of concentration Sex In a comparison of average inactive responses in the first 3 and last 3 sessions, animals in the THC group had higher inactive responding compared to the Veh group at both time points, but in the THC group, inactive responding decreased from the first to the last 3-session block (p<0.01). Inactive responding did not significantly decrease over time in Veh animals (p>0.05), perhaps due to a floor effect.

**Fig. 3.**
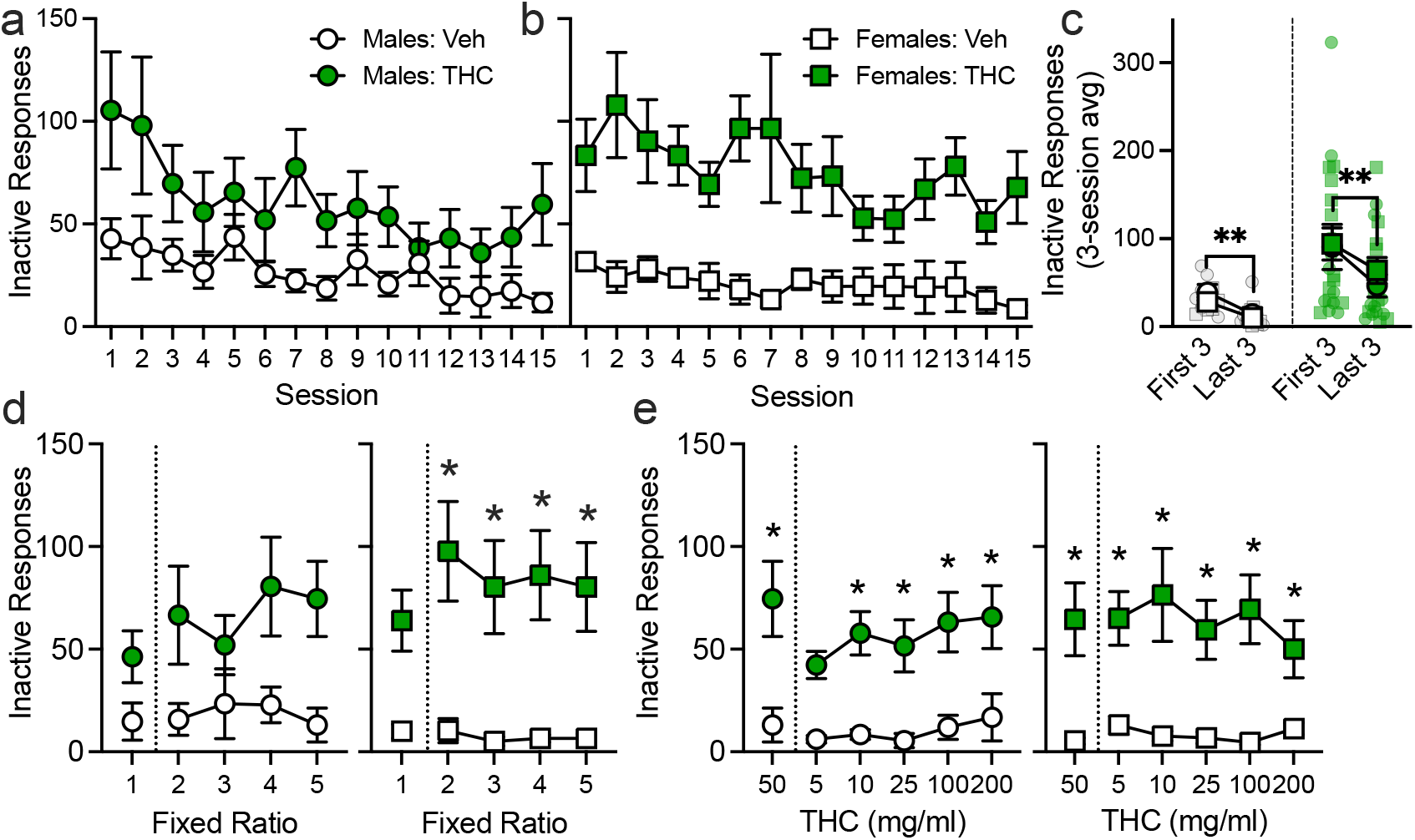
Inactive responses made by male (a) and female (b) rats in THC or Veh vapor groups an FR1 schedule. Responses decreased from the first 3 sessions to the last 3 sessions (c). Data are Mean± SEM; N=5-6/sex in vehicle group (white symbols), N=12/sex in THC group (green symbols). Males shown in circle symbols and females shown in square symbols. ^*^ denotes p<0.05 vs. Veh vapor group; # denotes a difference from FR1 or 50 mg/ml (training conditions)

In an analysis of inactive responses across FR schedules, there was a significant main effect of THC (F (1, 31) = 9.44, p<0.01), but not FR (F (3.497, 108.4) = 1.22, p=0.31) nor sex F (1, 31) = 0.03, p=0.85). While inactive responding was higher in both male and female rats in the THC group, post-hoc tests were significant only between females in the THC group vs. Veh group under all FRs (FR1, p=0.08; FR4-5, p’s>0.05). The higher inactive responding in THC males was not significantly different than the Veh group (p’s>0.05).

In the analysis of inactive responses while THC concentrations were varied, there was a significant main effect of THC (F (1, 31) = 13.22, p<0.01), but not concentration (F (2.54, 78.67) = 0.67), p=0.55) nor sex F (1, 31) = 0.01, p=0.92), and there were no significant interactions. Rats in the THC groups had higher inactive responding compared with Veh groups across varying THC concentrations.

### 3.5 Effect of pre-exposure condition on later responding for THC or Veh vapor

THC pre-exposure significantly increased self-administration of THC vapor. In an evaluation of THC deliveries obtained under FR1 schedule of reinforcement (Veh-THC: N=6 males, 6 females vs. THC-THC: N=12 males, 12 females), there were main effects of pre-exposure condition (F (1, 32) = 6.73, p<0.05) and session (F(5.56, 178.0) = 4.45, p<0.001) on deliveries (Fig. 4a-b). Pre-exposure condition also increased responding for THC vapor under increasing FR requirements (main effect of pre-exposure: F (1, 31) = 5.80, p<0.05) and as the THC concentration was varied (main effect of pre-exposure: F (1, 31) = 10.86, p<0.01; interaction of pre-exposure x FR: F(5, 155) = 4.03, p<0.01) (Fig. 4c-d).

**Fig. 4.**
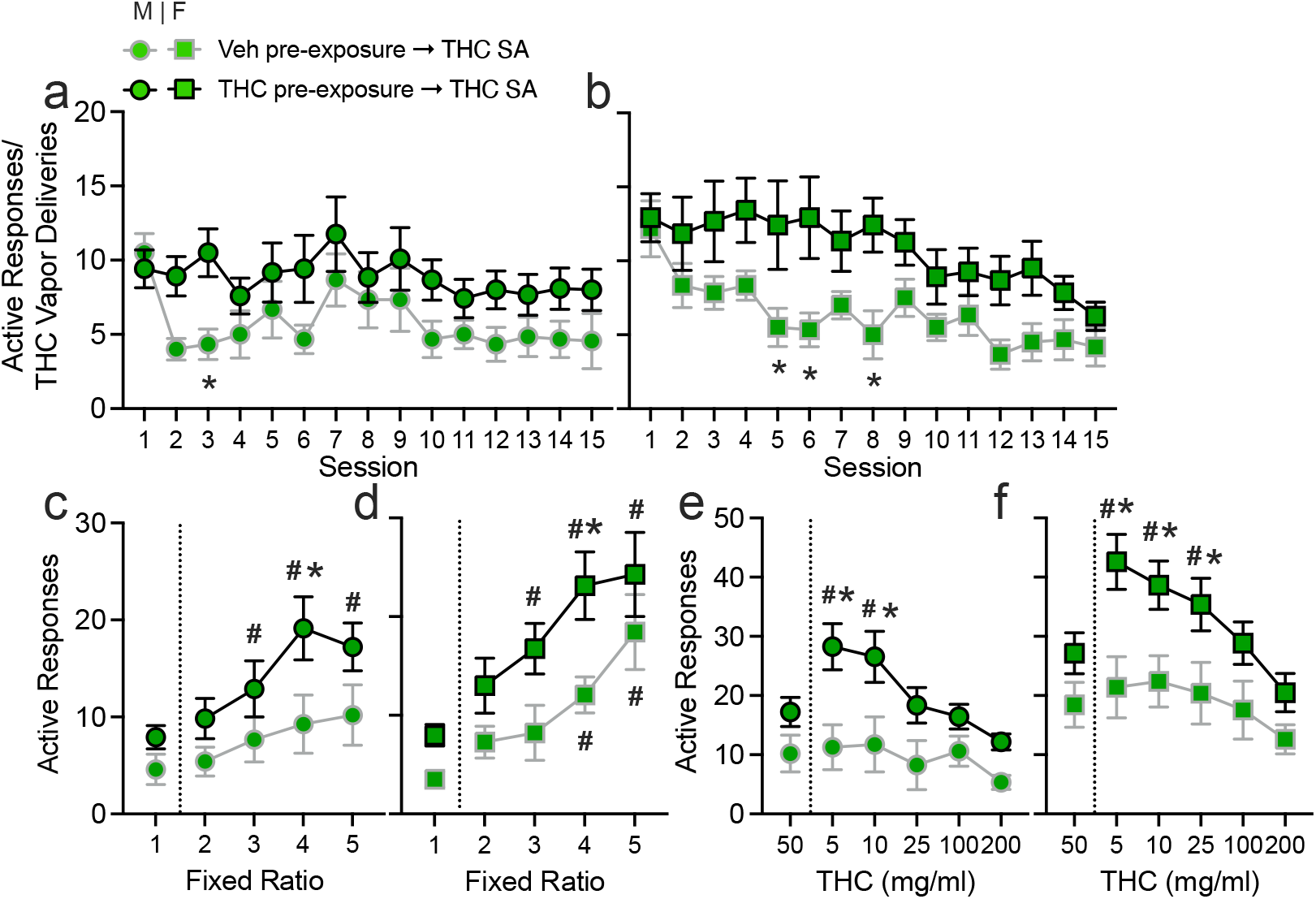
Effects of pre-exposure condition (THC or Veh vapor) on subsequent responding for THC vapor under an FR1 schedule (a-males; b-females), across FR1-5 schedules of reinforcement (c-males; d-females), and across varying THC concentrations (e-males; f-females). Symbols denote rats pre-exposed to THC (black outline) or Veh (gray outline) vapor; Males shown in circle symbols and females shown in square symbols. Data from THC-THC group (black outlines) are the same as shown in Fig 1. Data are Mean± SEM; N=12/sex/group. ^*^ denotes p<0.05 vs. Veh pre-exposure group; # denotes a difference from FR1 or 50 mg/ml (training conditions)

THC pre-exposure did not affect self-administration of Veh vapor (Fig. 5). Specifically, there was no effect of pre-exposure condition on Veh vapor deliveries under the FR1 schedule (F (1, 32) = 1.51, p=0.23), nor for responding for Veh vapor under increasing FR requirements (F (1, 31) = 0.12, p=0.73), nor when the THC concentration was varied (F (1, 31) = 0.14, p=0.71). There were also no interactions between pre-exposure and any other factors.

**Fig. 5.**
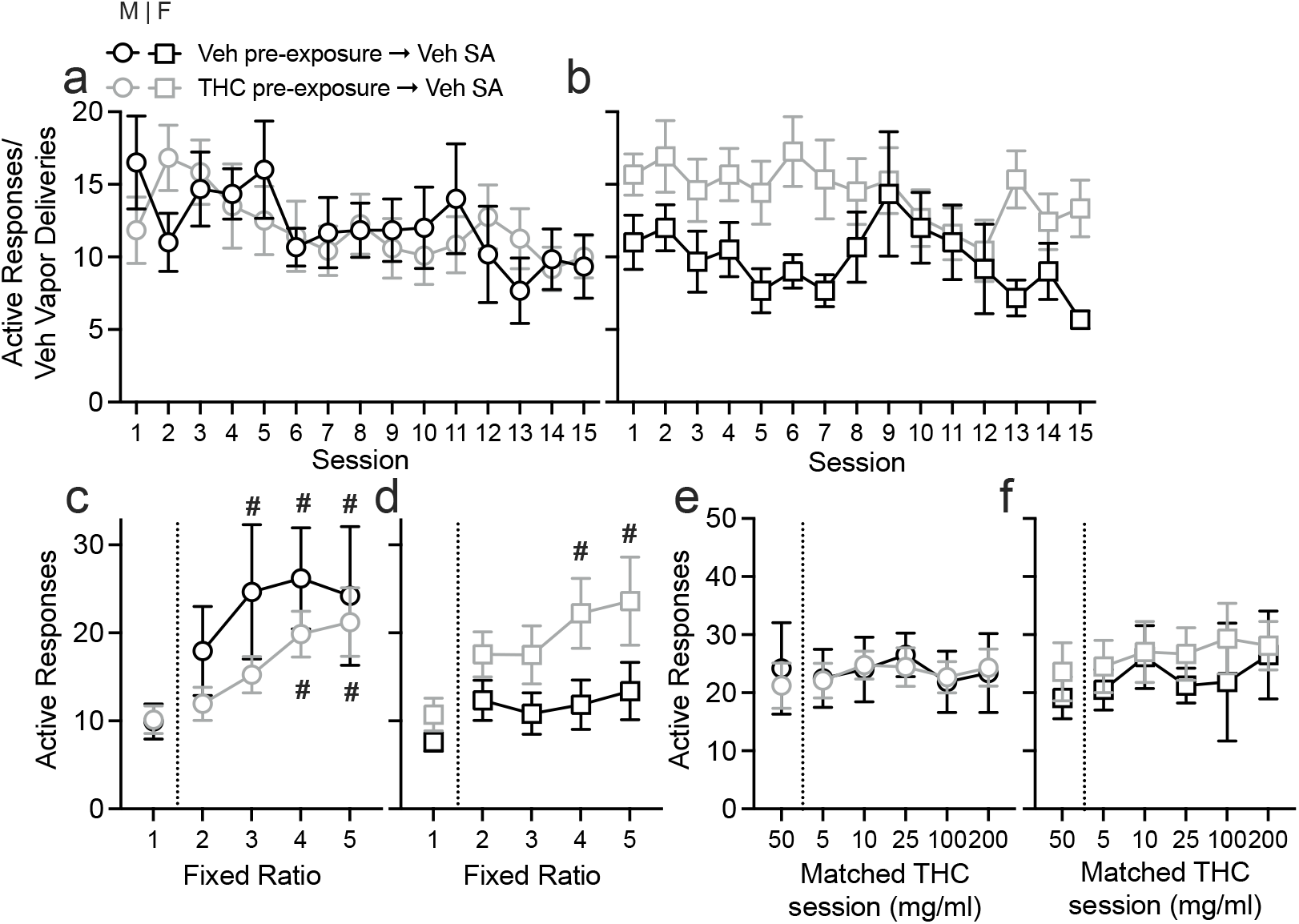
Effects of pre-exposure condition on responding for Veh vapor under an FR1 schedule (a-males; b-females), across FR1-5 schedules of reinforcement (c-males; d-females), and across varying THC concentrations (e-males; f-females). Symbols denote rats pre-exposed to THC (black outline) or Veh (gray outline) vapor; Males shown in circle symbols and females shown in square symbols. Data from Veh-Veh groups (black outlines) are the same as shown in Fig 1. Data are Mean± SEM; N=5-6/sex/group. # denotes a difference from FR1 or 50 mg/ml (training conditions)

## 4. Discussion

This study demonstrated that male and female rats will voluntarily administer THC vapor and will do so in a sustained manner across several months. Similar to previous studies (Freels et al. 2020; Glodosky et al. 2020; Freels et al. 2024), we observed no differences in self-administration of THC and Veh vapor under FR1 schedule of reinforcement in males and female rats. Rats in the THC group exerted more effort as the response cost per vapor delivery was increased, and also titrated their intake of THC in a predictable U-shaped pattern as the drug concentration was varied. THC vapor had reinforcing properties in female rats: female rats responding for THC vapor did so at higher levels than female rats responding for Veh vapor under FR4-5 schedules of reinforcement. Interestingly, males responding for the Veh vapor had higher intake compared to males in the THC vapor group, and also increased their responding as the FR increased. Female rats in the Veh vapor group had no change in their responding as the FR increased. Results showing lower responding for THC vapor compared to Veh vapor in male rats are in contrast to Freels et al. (2020) where there was higher self-administration of vaporized THC-rich cannabis extract in male rats above vaporized Veh. This study differs from Freels et al. (2020) in several key ways: we used isolated THC mixed in 100% propylene glycol (PG), whereas the prior study used cannabis extracts and a vehicle solution of 80% propylene glycol and 20% vegetable glycerin. While our THC training dose (50 mg/ml) was matched to this prior study, these results indicate that there may be differences in 1) reinforcing properties of whole plant extracts that contain a variety of cannabinoids and terpenes vs. THC alone and/or 2) potentially reinforcing effects of the PG vehicle vapor solution. It has been shown that people are using drug-free vapes (Tokle et al. 2022), suggesting that these e-liquids do have reinforcing properties on their own, warranting further study.

Next in this study, we evaluated responding for vapor across various concentrations of THC to produce a dose effect curve. In both males and females, we had higher levels of responding for lower THC concentrations (5-25 mg/ml) compared to the training dose of 50 mg/ml. In females, these lower concentrations also produced higher levels of responding compared to the Veh vapor group. This aligns with our understanding that higher doses of THC may be aversive and demonstrate that lower doses may engender higher self-administration. Future studies will be aimed at investigating the use of lower concentrations of THC during training.

Using our experimental setup, we saw very high levels of inactive nose poke responding in THC self-administering animals specifically. One potential reason may be that these animals were trained on food self-administration in operant chambers where a nose poke response on a key resulted in delivery of a food pellet into a hopper in the center. The response operanda in the vapor self-administration chambers is a nose poke or ‘entry’ into a hole. Its possible that there was generalization of the food hopper with the nose poke hole, and this behavior was related to food-seeking. If this were true, we would expect to see extinction in this behavior over time, and while rats in the Veh vapor group did decrease inactive responses to low levels, the THC vapor group’s inactive responding remained very high throughout the months of testing. Another explanation is that the higher inactive responding in the THC vapor group is due to locomotor stimulating effects of the THC. In our prior study (Moore et al. 2024), we observed that THC vapor was stimulating in female rats, and this has also been observed at lower doses of THC administration (Sanudo-Pena et al. 2000; Katsidoni et al. 2013). Based on responding patterns of rats in the THC group to titrate intake, including increasing their responding while the drug concentration was reduced (e.g., 5 mg/ml), we do not believe there was a failure to discriminate between the active and inactive nosepoke.

When comparing self-administration behaviors of animals that had either been pre-exposed to THC or not (i.e., Veh pre-exposure), we saw that pre-exposure to THC resulted in subsequently higher responding for THC vapor that was sustained over time. Most interestingly, on the first day of vapor self-administration, there was no difference in number of responses made, but responding of rats who were THC-naïve prior to day 1 progressively declined across the following sessions. In males, this effect was very pronounced, with a sharp decline in responding immediately in the next session, while in females this decline was gradual and most notable after session 5. THC naive animals therefore decreased their intake over time across FR1 sessions, had very small increases in responding as the FR increased, and showed a very shallow dose-effect curve for THC. There was no effect of pre-exposure condition on self-administration of vehicle vapor.

This is a promising model of voluntary cannabis use, replicating what has been shown with vaporized cannabis extracts. We show that pre-exposure is critical for engendering higher intake/ intake above vehicle levels in females. Future studies should examine initiation of self-administration using even lower dose concentrations than the 50 mg/ml training condition in the current study, and examine other factors that may influence vehicle responding (e-liquid used, timeouts between vapor deliveries, response schedules and session duration etc. It is important to understand the reinforcing effects of individual cannabis components, including THC, on their own. Using vapor self-administration procedures we can assess which cannabis constituents have reinforcing effects, and/or interact with THC to alter its reinforcing effects.

## BIBLIOGRAPHY

Arroyo M, Markou A, Robbins TW, and Everitt BJ. (1998) Acquisition, maintenance and reinstatement of intravenous cocaine self-administration under a second-order schedule of reinforcement in rats: effects of conditioned cues and continuous access to cocaine. Psychopharmacology (Berl), 140: 331–44. 10.1007/s002130050774

Budney AJ, Sargent JD, and Lee DC. (2015) Vaping cannabis (marijuana): parallel concerns to e-cigs? Addiction, 110: 1699–704. 10.1111/add.13036

Calipari ES, Siciliano CA, Zimmer BA, and Jones SR. (2015) Brief intermittent cocaine self-administration and abstinence sensitizes cocaine effects on the dopamine transporter and increases drug seeking. Neuropsychopharmacology, 40: 728–35. 10.1038/npp.2014.238

Carnicella S, Ron D, and Barak S. (2014) Intermittent ethanol access schedule in rats as a preclinical model of alcohol abuse. Alcohol, 48: 243–52. 10.1016/j.alcohol.2014.01.006

Deiana S, Fattore L, Spano MS, Cossu G, Porcu E, Fadda P, and Fratta W. (2007) Strain and schedule-dependent differences in the acquisition, maintenance and extinction of intravenous cannabinoid self-administration in rats. Neuropharmacology, 52: 646–54. 10.1016/j.neuropharm.2006.09.007

Deroche-Gamonet V. (2020) The relevance of animal models of addiction. Addiction, 115: 16–17. 10.1111/add.14821

Fattore L, Cossu G, Martellotta CM, and Fratta W. (2001) Intravenous self-administration of the cannabinoid CB1 receptor agonist WIN 55,212-2 in rats. Psychopharmacology (Berl), 156: 410–6. 10.1007/s002130100734

Fattore L, Fadda P, and Fratta W. (2007) Endocannabinoid regulation of relapse mechanisms. Pharmacol. Res., 56: 418–27. 10.1016/j.phrs.2007.09.004

Fattore L, Spano MS, Altea S, Angius F, Fadda P, and Fratta W. (2007) Cannabinoid self-administration in rats: sex differences and the influence of ovarian function. Br. J. Pharmacol., 152: 795–804. 10.1038/sj.bjp.0707465

Freels TG, Baxter-Potter LN, Lugo JM, Glodosky NC, Wright HR, Baglot SL, Petrie GN, Yu Z, Clowers BH, Cuttler C, Fuchs RA, Hill MN, and McLaughlin RJ. (2020) Vaporized Cannabis Extracts Have Reinforcing Properties and Support Conditioned Drug-Seeking Behavior in Rats. J. Neurosci., 40: 1897–908. 10.1523/JNEUROSCI.2416-19.2020

Freels TG, Westbrook SR, Zamberletti E, Kuyat JR, Wright HR, Malena AN, Melville MW, Brown AM, Glodosky NC, Ginder DE, Klappenbach CM, Delevich KM, Rubino T, and McLaughlin RJ. (2024) Sex Differences in Response-Contingent Cannabis Vapor Administration During Adolescence Mediate Enduring Effects on Behavioral Flexibility and Prefrontal Microglia Activation in Rats. Cannabis Cannabinoid Res, 9: e1184–e96. 10.1089/can.2023.0014

Giroud C, de Cesare M, Berthet A, Varlet V, Concha-Lozano N, and Favrat B. (2015) E-Cigarettes: A Review of New Trends in Cannabis Use. Int J Environ Res Public Health, 12: 9988–10008. 10.3390/ijerph120809988

Glodosky NC, Cuttler C, Freels TG, Wright HR, Rojas MJ, Baglot SL, Hill MN, and McLaughlin RJ. (2020) Cannabis vapor self-administration elicits sex- and dose-specific alterations in stress reactivity in rats. Neurobiol Stress, 13: 100260. 10.1016/j.ynstr.2020.100260

Goodwin AK. (2016) An intravenous self-administration procedure for assessing the reinforcing effects of hallucinogens in nonhuman primates. J. Pharmacol. Toxicol. Methods, 82: 31–36. 10.1016/j.vascn.2016.07.004

Holgate JY, Shariff M, Mu EW, and Bartlett S. (2017) A Rat Drinking in the Dark Model for Studying Ethanol and Sucrose Consumption. Front Behav Neurosci, 11: 29. 10.3389/fnbeh.2017.00029

Justinova Z, Munzar P, Panlilio LV, Yasar S, Redhi GH, Tanda G, and Goldberg SR. (2008) Blockade of THC-seeking behavior and relapse in monkeys by the cannabinoid CB(1)-receptor antagonist rimonabant. Neuropsychopharmacology, 33: 2870–7. 10.1038/npp.2008.21

Justinova Z, Tanda G, Redhi GH, and Goldberg SR. (2003) Self-administration of delta9-tetrahydrocannabinol (THC) by drug naive squirrel monkeys. Psychopharmacology (Berl), 169: 135–40. 10.1007/s00213-003-1484-0

Katsidoni V, Kastellakis A, and Panagis G. (2013) Biphasic effects of Delta9-tetrahydrocannabinol on brain stimulation reward and motor activity. Int J Neuropsychopharmacol, 16: 2273–84. 10.1017/S1461145713000709

Kawa AB, Bentzley BS, and Robinson TE. (2016) Less is more: prolonged intermittent access cocaine self-administration produces incentive-sensitization and addiction-like behavior. Psychopharmacology (Berl), 233: 3587–602. 10.1007/s00213-016-4393-8

Kenny PJ, Chen SA, Kitamura O, Markou A, and Koob GF. (2006) Conditioned withdrawal drives heroin consumption and decreases reward sensitivity. J. Neurosci., 26: 5894–900. 10.1523/JNEUROSCI.0740-06.2006

Lefever TW, Marusich JA, Antonazzo KR, and Wiley JL. (2014) Evaluation of WIN 55,212-2 self-administration in rats as a potential cannabinoid abuse liability model. Pharmacol. Biochem. Behav., 118: 30–5. 10.1016/j.pbb.2014.01.002

Mandyam CD, Wee S, Crawford EF, Eisch AJ, Richardson HN, and Koob GF. (2008) Varied access to intravenous methamphetamine self-administration differentially alters adult hippocampal neurogenesis. Biol Psychiatry, 64: 958–65. 10.1016/j.biopsych.2008.04.010

Markou A, Li J, Tse K, and Li X. (2016) Cue-induced nicotine-seeking behavior after withdrawal with or without extinction in rats. Addict. Biol. 10.1111/adb.12480

Markou A, Weiss F, Gold LH, Caine SB, Schulteis G, and Koob GF. (1993) Animal models of drug craving. Psychopharmacology (Berl), 112: 163–82. 10.1007/BF02244907

Martellotta MC, Cossu G, Fattore L, Gessa GL, and Fratta W. (1998) Self-administration of the cannabinoid receptor agonist WIN 55,212-2 in drug-naive mice. Neuroscience, 85: 327–30. 10.1016/S0306-4522(98)00052-9

Moore CF, Davis CM, Sempio C, Klawitter J, Christians U, and Weerts EM. (2024) Delta(9)-Tetrahydrocannabinol Vapor Exposure Produces Conditioned Place Preference in Male and Female Rats. Cannabis Cannabinoid Res, 9: 111–20. 10.1089/can.2022.0175

Morean ME, Lipshie N, Josephson M, and Foster D. (2017) Predictors of Adult E-Cigarette Users Vaporizing Cannabis Using E-Cigarettes and Vape-Pens. Subst. Use Misuse, 52: 974–81. 10.1080/10826084.2016.1268162

Murray JE, and Bevins RA. (2010) Cannabinoid conditioned reward and aversion: behavioral and neural processes. ACS Chem Neurosci, 1: 265–78. 10.1021/cn100005p

O’Dell LE, and Koob GF. (2007) ’Nicotine deprivation effect’ in rats with intermittent 23-hour access to intravenous nicotine self-administration. Pharmacol. Biochem. Behav., 86: 346–53. 10.1016/j.pbb.2007.01.004

Panlilio LV, Justinova Z, and Goldberg SR. (2010) Animal models of cannabinoid reward. Br. J. Pharmacol., 160: 499–510. 10.1111/j.1476-5381.2010.00775.x

Passarotti A, Crane NA, Hedeker D, and Mermelstein RJ. (2015) Longitudinal trajectories of marijuana use from adolescence to young adulthood. Addict. Behav., 45: 301–08. 10.1016/j.addbeh.2015.02.008

Sanudo-Pena MC, Romero J, Seale GE, Fernandez-Ruiz JJ, and Walker JM. (2000) Activational role of cannabinoids on movement. Eur. J. Pharmacol., 391: 269–74. 10.1016/s0014-2999(00)00044-3

Spindle TR, Cone EJ, Goffi E, Weerts EM, Mitchell JM, Winecker RE, Bigelow GE, Flegel RR, and Vandrey R. (2020) Pharmacodynamic effects of vaporized and oral cannabidiol (CBD) and vaporized CBD-dominant cannabis in infrequent cannabis users. Drug Alcohol Depend., 211: 107937. 10.1016/j.drugalcdep.2020.107937

Takahashi RN, and Singer G. (1979) Self-administration of delta 9-tetrahydrocannabinol by rats. Pharmacol. Biochem. Behav., 11: 737–40. 10.1016/0091-3057(79)90274-0

Takahashi RN, and Singer G. (1980) Effects of body weight levels on cannabis self-injection. Pharmacol. Biochem. Behav., 13: 877–81. 10.1016/0091-3057(80)90222-1

Tanda G. (2016) Preclinical studies on the reinforcing effects of cannabinoids. A tribute to the scientific research of Dr. Steve Goldberg. Psychopharmacology (Berl), 233: 1845–66. 10.1007/s00213-016-4244-7

Tanda G, Munzar P, and Goldberg SR. (2000) Self-administration behavior is maintained by the psychoactive ingredient of marijuana in squirrel monkeys. Nat. Neurosci., 3: 1073–4. 10.1038/80577

Tokle R, Brunborg GS, and Vedøy TF. (2022) Adolescents’ use of nicotine-free and nicotine e-cigarettes: A longitudinal study of vaping transitions and vaper characteristics. Nicotine Tob Res, 24: 400–07. 10.1093/ntr/ntab192

van Ree JM, Slangen JL, and de Wied D. (1978) Intravenous self-administration of drugs in rats. J. Pharmacol. Exp. Ther., 204: 547–57. 10.1016/S0022-3565(25)31172-9

Wikler A. (1973) Dynamics of drug dependence: implications of a conditioning theory for research and treatment. Arch Gen Psychiatry, 28: 611–16. 10.1001/archpsyc.1973.01750350005001

